# Multiplexed detection of bacterial nucleic acids using Cas13 in droplet microarrays

**DOI:** 10.1101/2021.11.12.468388

**Authors:** Sri Gowtham Thakku, Cheri M. Ackerman, Cameron Myhrvold, Roby P. Bhattacharyya, Jonathan Livny, Peijun Ma, Giselle Isabella Gomez, Pardis C. Sabeti, Paul C. Blainey, Deborah T. Hung

## Abstract

Rapid and accurate diagnosis of infections is fundamental to individual patient care and public health management. Nucleic acid detection methods are critical to this effort, but are limited either in the breadth of pathogens targeted or by the expertise and infrastructure required. We present here a high-throughput system that enables rapid identification of bacterial pathogens, bCARMEN, which utilizes: (1) modular CRISPR-Cas13-based nucleic acid detection with enhanced sensitivity and specificity; and (2) a droplet microfluidic system that enables thousands of simultaneous, spatially multiplexed detection reactions at nanoliter volumes; and (3) a novel pre-amplification strategy that further enhances sensitivity and specificity. We demonstrate bCARMEN is capable of detecting and discriminating 52 clinically relevant bacterial species and several key antibiotic resistance genes. We further develop a proof of principle system for use with stabilized reagents and a simple workflow with optical readout using a cell phone camera, opening up the possibility of a rapid point-of-care multiplexed bacterial pathogen identification and antibiotic susceptibility testing.

**Significance Statement:** In this paper, we use a novel primer design method combined with droplet-based CRISPR Cas13 detection to distinguish 52 clinically relevant bacterial pathogens in a single assay. We also apply the method to detect and distinguish a panel of major antibiotic resistance genes, which is of critical importance in this era of rising antibiotic resistance. Finally, we make key advances towards making our diagnostic assay deployable at the point-of-care, with a simplified emulsion-free assay process that uses mobile phone camera for detection and reduces infrastructure/skilled labor requirements.

## Introduction

Infections represent a substantial fraction of worldwide disease burden^1^, and their rapid detection is critical to both patient care and containment. Diagnosis of bacterial infections has long relied on culture followed by biochemical assays^2^, which can take days to return an answer and which requires significant laboratory infrastructure, or mass spectrometry^3^. Molecular diagnostic tools have also begun to see increasing use in clinical practice with the advantages of high sensitivity, specificity, and speed^2,4^. Nucleic acid amplification tests (NAATs) in particular enable highly specific targeting of genomic regions, thereby allowing greater levels of taxonomic resolution. Such diagnostic approaches have been critical to the containment of the ongoing COVID-19 pandemic.^5,6^ In recent years, a number of CRISPR-based NAAT assays have emerged, including SHERLOCK^7–9^ (specific high-sensitivity enzymatic reporter unlocking) and DETECTR^10^ (DNA endonuclease-targeted CRISPR trans reporter), which employ the CRISPR effectors Cas13 and Cas12, respectively. Recently, SHERLOCK received approval for clinical use for the detection of SARS-CoV-2^11^. CRISPR effector-based NAAT have two amplification stages that each impose distinct specificity requirements, thereby ensuring good specificity as well as sensitivity. The first step is pre-amplification with a method such as PCR (polymerase chain reaction), RPA (recombinase polymerase amplification) or LAMP (loop-mediated amplification). The second step is target-sequence dependent amplified signal generation by one of the CRISPR effector systems. These components can be viewed as a modular NAAT tool kit that can be deployed in new configurations that expand testing possibilities^12^.

An important limitation of current NAATs using, for example, RPA, LAMP (loop-mediated isothermal amplification), or current CRISPR-based technologies is the need for a diagnostic hypothesis to guide targeted inquiry. Testing a broader panel of pathogens is needed to enable more comprehensive testing in the absence of a clear diagnostic hypothesis. Meanwhile, resistance gene detection would provide additional information to support treatment selection and epidemiological tracking. Whereas small panel tests have been developed for PCR-based NAATs^13^, the constraints of highly multiplexed amplification and barcoding limit the size of these panels to tens of pathogen targets.^14^ In contrast, next-generation sequencing (NGS) offers unbiased identification of pathogens^15^ with increasingly accurate prediction of antibiotic resistance phenotypes.^16,17^ However, NGS assays require considerable infrastructure and remain time-consuming, expensive, and complex to interpret. An ideal diagnostic assay would combine the sensitivity, specificity, and speed of NAATs with the breadth of pathogen identification offered by sequencing in a format that requires minimal infrastructure.

We have previously described Combinatorial Arrayed Reactions for Multiplexed Evaluation of Nucleic acids (CARMEN) and demonstrated its application to the detection of a large panel of human-associated viruses^18^. CARMEN combines the modularity of CRISPR-based nucleic acid sensing with the throughput capabilities of the DropArray platform, a miniaturized microwell system we developed to enable comprehensive, high throughput combinatorial experiments.^19,20^ CARMEN encapsulates pre-amplified nucleic acid targets and Cas13-guided detection sets into distinct nanoliter droplets in order to run tens to hundreds of thousands of pairs of target-guide detection reactions in parallel (Figure 1). Here we employ a novel assay design strategy and showcase the use of CARMEN for the discrimination of bacterial species and detection of antibiotic susceptibility genes, and call it bCARMEN (bacterial CARMEN.)

**Figure 1:**
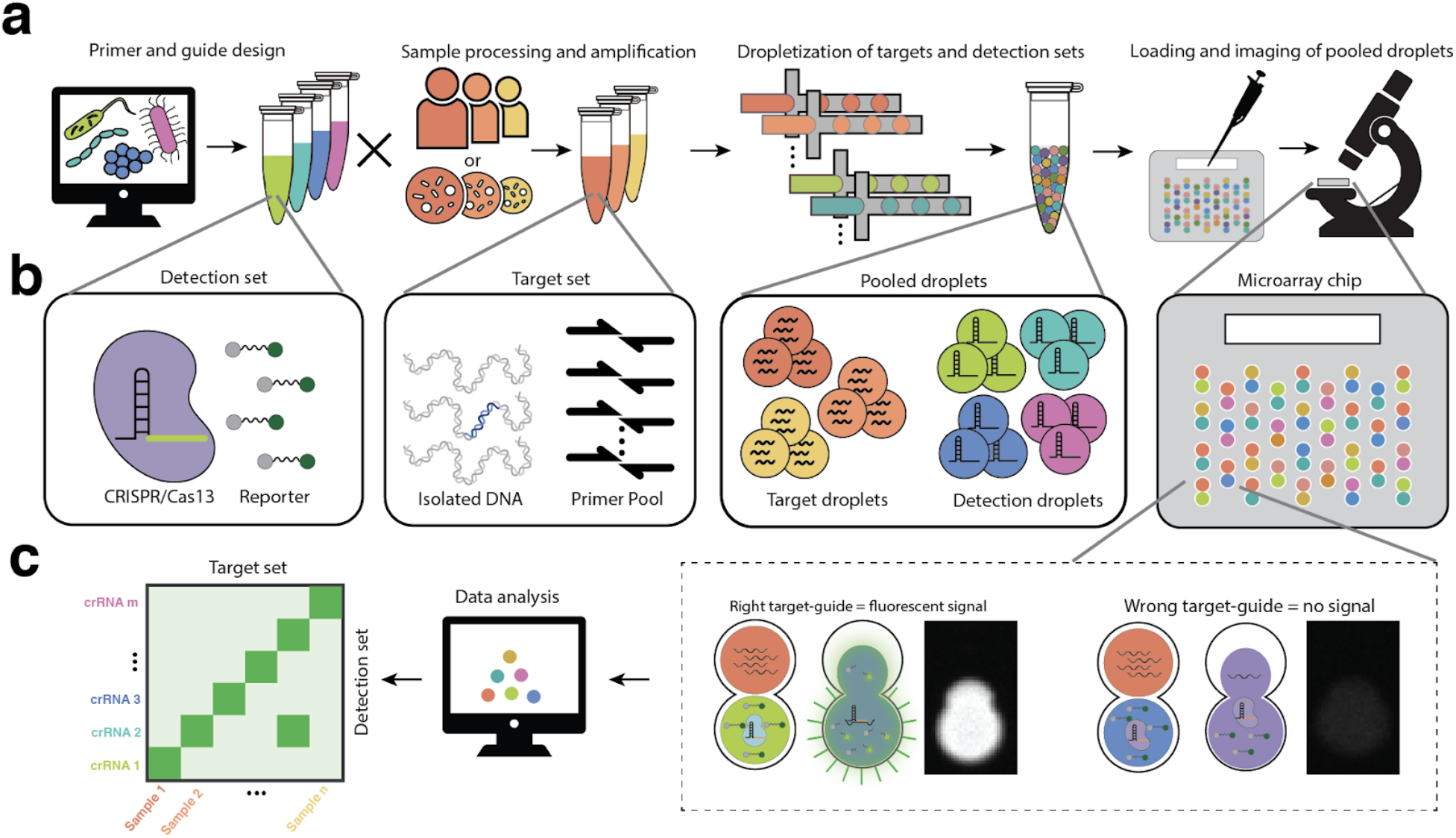
bCARMEN-Cas13 workflow from assay design to readout. **a**, bCARMEN workflow, which includes primer and guide design, sample processing and amplification, dropletization of targets and detection sets, loading and imaging of pooled droplets **b**, Components of items in workflow; detection set contains Cas13 protein, crRNA, reporter, and other reagents; target set includes DNA from sample, primer pool and PCR reagents; pooled droplets include all target and detection droplets; each microarray chip has wells that hold two droplets, and stochastic loading ensures every pairwise combination is represented **c**, Data analysis workflow; wells with the right target-guide pair show strong fluorescence signal while wells with pairs show no signal; data analysis of images results in a heatmap for easy interpretation of results.

Bacterial genomes are far more complex than viral genomes and include highly conserved and diverse regions.^21^ These features present unique opportunities to leverage both the conserved regions for amplification across a broad panel of bacterial species as well as the unique regions to distinguish individual species. While the 16s ribosomal gene might seem ideal for this purpose and has indeed been used extensively for taxonomic classification,^22,23^ we found that it provided insufficient resolution to distinguish relevant bacterial pathogens. We thus turned to a set of housekeeping genes present in most bacterial species^24^. We identified regions in these genes that are highly variable across species, allowing their unique identification, but are flanked by highly conserved regions to enable their collective amplification. We found that regions of the *topA* gene fulfilled these criteria and served as a good target to enable identification and differentiation of 52 clinically relevant species.

Clinical management of bacterial infections requires not only species identification but also antibiotic susceptibility testing (AST), particularly in this era of rising antibiotic resistance. AST informs treatment regimens and enables the tracking of drug resistance across geographies and time.^25^ Current AST methods commonly involve exposure of clinical isolates to drugs and bacterial growth as the readout.^2^ In certain cases, genotypic resistance markers are clearly predictive of susceptibility to high-value antibiotics. Thus, we expanded bCARMEN’s application to detecting common bacterial resistance genes in clinical isolates, including genes conferring resistance to methicillin in Staphylococcus *aureus*, vancomycin in Enterococcus *faecalis and E. faecium*, and carbapenems in different Enterobacteriaceae species.

Finally, important requirements for wide adoption of a diagnostic assay are speed and ease of use. We addressed these challenges by showing proof-of-concept for a droplet-free workflow that includes cell phone imaging for readout. This streamlined approach significantly reduced time to result and infrastructure requirements and highlights the potential for broad deployment.

## Results

### Primer and guide design for bacterial species-specific detection

We set out to apply the CRISPR effector NAAT toolset to design a broad bacterial species identification panel. Our goals were to achieve (1) broad coverage by the amplification primers to capture all species of interest and (2) high specificity in the crRNA guides to identify and distinguish between species of interest (Figure 2a). Unlike common NAAT approaches where a single reaction step must achieve all the amplification required for sensitivity while maintaining specificity for targets to be differentiated, CRISPR effector-based NAAT like SHERLOCK can carry out robust pre-amplification with relaxed specificity while relying on the CRISPR detection step for additional specificity.

**Figure 2:**
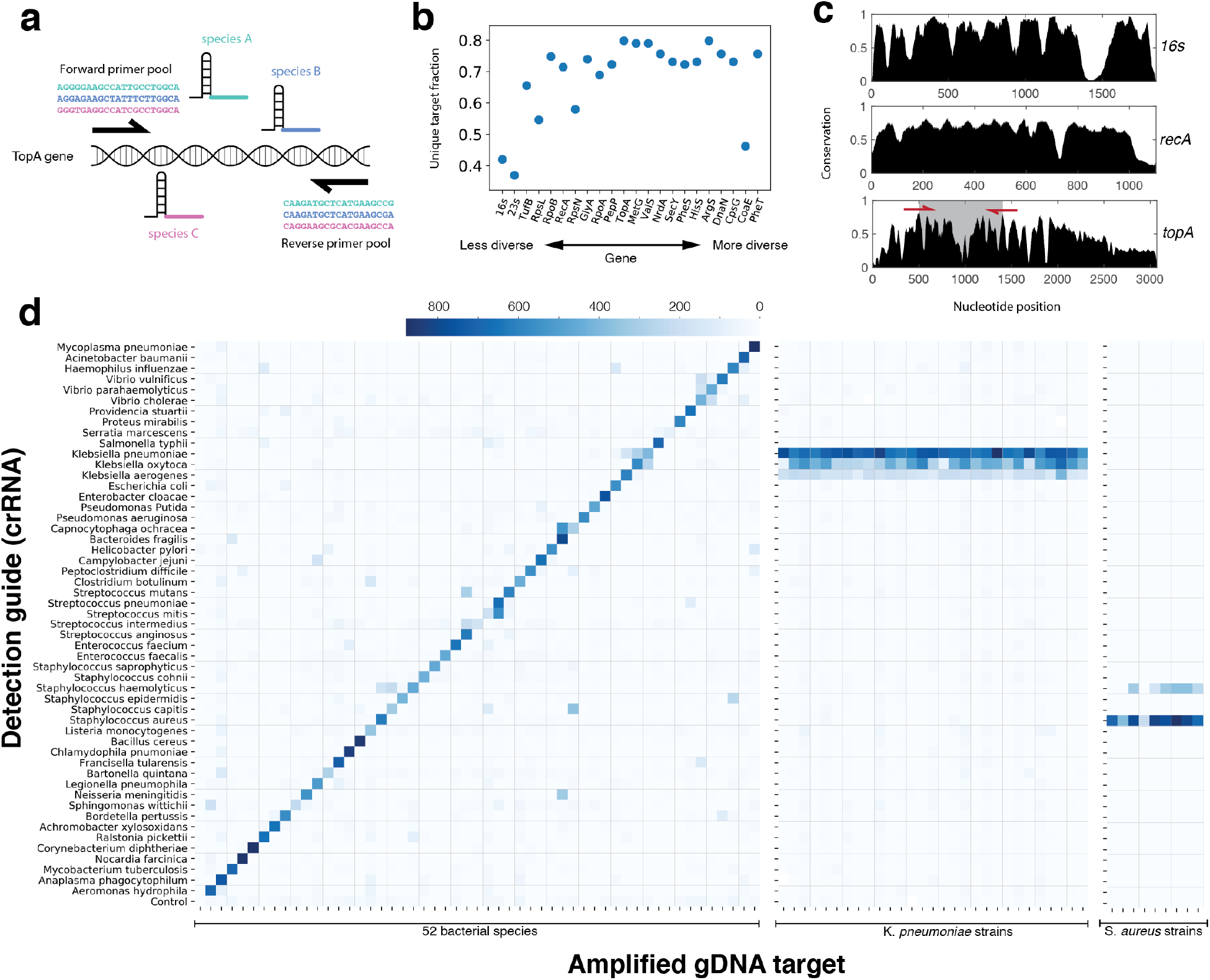
Bacterial species identification using bCARMEN-Cas13. **a**, Schematic of *topA* gene amplicon with forward and reverse primer pools and species specific guide binding site (each color refers to a distinct species). **b**, Fraction of uniquely identifiable species for 20 housekeeping genes. **c**, Conservation as a function nucleotide position across 52 bacterial species plotted for 16s, *recA* and *topA* genes. **d**, Testing a large bacterial panel using bCARMEN-Cas13.Detection sets containing species specific crRNA guides are along the vertical axis. Bacterial gDNA amplified using a one-pot pooled PCR is along the horizontal axis (order of 52 species is the same as detection set guides).

We first identified 52 bacterial pathogens that cause some of the most commonly reported bacterial infections (Table S1). We started by testing 16s universal primers because they are commonly used for amplification followed by targeted sequencing for taxonomic identification; however, we found that the 16S rRNA gene is not sufficiently diverse to support species-level discrimination by the second, crRNA detection step (Figure S2). To expand our search, we analyzed other conserved bacterial genes. We estimated the degree of conservation of these genes across species and plotted this as a function of nucleotide position and identified candidate genes with partially conserved regions flanking those of high diversity (Figure 2b, 2c, S1). We wanted the region between forward and reverse primers to span crRNA targets (28 nucleotides) with sufficient diversity to enable species differentiation, while keeping the total amplicon length short enough to enable efficient PCR amplification (<1000 nucleotides). Primers targeting the conserved segments provide broad coverage for amplification, while crRNAs targeting the intervening diverse regions enable high-resolution discrimination of related species (Figure 2a). We looked for genes which had regions with at least 80% conservation flanking regions with less than 50% conservation. After shortlisting three genes (valS, topA, rpoB), we found that the *topA* gene, which encodes the DNA topoisomerase 1 protein, offers the desired advantageous juxtaposition of sequence diversity and conservation. To enable discrimination of a large bacterial panel, we chose an amplicon (~1000 bases) encompassing multiple variable regions able to accommodate crRNA targets unique to each of the 52 species. We curated a gene sequence database from the identified list of 52 pathogenic bacterial species (median of 10 strains per species) and used it as the basis to design amplification primers specific to each species with the maximum coverage of the strains representing each species. Since primers targeted regions of high but not perfect conservation (>80%, <100%), sufficient sequence variation exists that distinct primer sequences were required for each species. While multiplexing PCR beyond a handful of primer sets is typically challenging, because we targeted the same homologous *topA* sequence in each species, the 52 primer pairs had a high degree of similarity, thereby enabling their multiplexing in a single PCR reaction.

Using the resulting amplicon sequences, we then applied ADAPT^26^ to design crRNA guides that optimally distinguish each species from all others. These guides were predicted not to crossreact (>3 base pair mismatches) with amplicons corresponding to all included strains from the 51 non-target pathogens in the panel. Additionally, a BLAST search was performed to ensure no predicted cross-reactivity with bacterial and human genomes outside our panel. We tested the 52 amplification primer sets and guides using genomic DNA extracted from the corresponding 52 bacterial species. These species included a mix of lab strains and clinical isolates and represented a more realistic test condition than short synthetic targets (Table S1).

After two rounds of design and testing, we found a set of 52 primer pairs that amplified their corresponding *topA* targets in 52/52 species with 48/52 (92%) crRNAs showing a significant signal above background (signal > 6 standard deviations above background). 23/52 crRNAs showed some cross-reactivity against a single additional target, and no crRNAs had more than one cross reactive signal (Figure S3). Even in the crRNAs that did show cross reactivity, the reactivity pattern across all crRNAs still uniquely identified the target species. To quantify this, we compared the expected reactivity pattern against the observed pattern and found a mean AUC of >0.99 across all guides (Figure S3). A mean of 10.8 replicates (wells containing a given crRNA-target pair) were generated in a single assay run, and 3 replicates were sufficient to make a call with >99% confidence (Figure S4.)

Since strains within a species can have single nucleotide polymorphisms (SNPs), we sought to demonstrate robustness of species identification across a larger number of strains of the same species. We tested 10 different strains of *Staphylococcus aureus* and 30 strains of *Klebsiella pneumoniae* with their respective primers and crRNAs. We found that the crRNAs for *S. aureus* and *K. pneumoniae* gave a strong positive signal for all their respective target strains (10/10 S. *aureus*, 30/30 K. *pneumoniae*). 7/10 S. *aureus* strains showed weak cross reactivity with the S. *haemolyticus* crRNA, while 27/30 K. *pneumoniae* strains showed weak cross reactivity with the *K. oxytoca* crRNA and 29/30 strains showed very weak cross reactivity with the *K. aerogenes* crRNA. In all cases, the signal from the target species crRNA was greater than that from the cross-reacting species crRNA.

### Detection of clinically relevant bacterial resistance genes

A key requirement of bacterial diagnostics in the clinic is antibiotic susceptibility testing (AST). To demonstrate the potential for bacterial CARMEN to detect genotypic resistance markers, we designed primers and guides for 14 different resistance genes spanning three classes of critical drug-resistant pathogens which the United States Centers for Disease Control has highlighted as serious or urgent threats: carbapenem-resistant Enterobacteriaceae (CRE), vancomycin-resistant enterococci (VRE) and methicillin-resistant *Staphylococcus aureus* (MRSA).^27^ We curated a database of resistant gene variants and designed primers to target conserved regions of each gene for amplification and detection. In order to showcase multiple possible assay designs, we applied different strategies for resistance gene detection (Figure 3a). This was also partly determined by the degree of conservation within members of each resistance gene family. In one strategy, we targeted a single gene with a unique primer pair and guide (*mecA*, *mecC*, *bla*_NDM-1_, *bla*_CTX-M-15_, *mcr1*). In a second strategy, exemplified by the detection of *bla*_KPC_, *bla*_VIM_, *bla*_IMP_, and o*xa48*-like genes, we collapsed the diversity of gene variants using a single crRNA probe targeting a well-conserved region. In a third strategy, we discriminated between different genes conferring a similar phenotypic resistance profile, by designing primers targeting conserved regions and crRNAs targeting more diverse regions, thus enabling epidemiological tracking of the spread of infectious pathogens or resistance elements. This third strategy was similar to that used in the bacterial identification panel, and we exemplified this strategy by designing crRNAs to detect a set of vancomycin resistance conferring (van) genes in enterococci. Primers and crRNAs for each resistance element were first tested using synthetic targets; we found that 14/14 crRNAs were selective for their target (Figure 3c, S4.) We then tested the assay using 26 clinical isolates with known genotypes. The assay detected 27/27 (100%) resistance genes (signal > 6 s.d. above background) and showed no off-target reactivity (no other signal > 6 s.d. above background).

**Figure 3:**
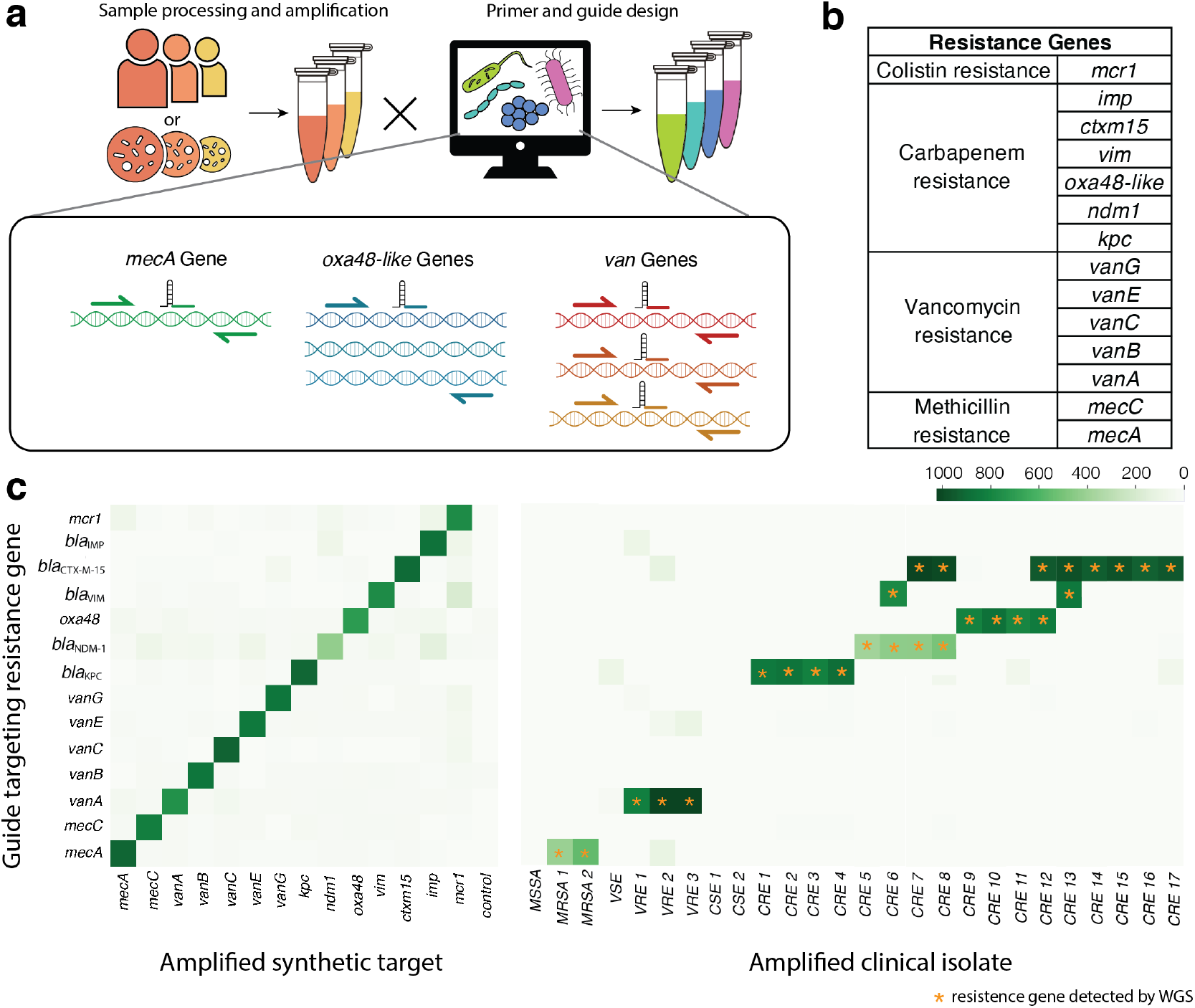
Bacterial resistance gene detection using bCARMEN-Cas13. **a**, Schematic showing the range of design strategies enabled by bCARMEN-Cas13. Variants of a class of genes can be discriminated (*van*, *mec*) or combined in a single detection (oxa48). **b**, List of bacterial resistance genes detected using bCARMEN-Cas13. **c**, Detection of synthetic targets and resistance genes in clinical isolates (MSSA = methicillin susceptible *Staphylococcus aureus*; MRSA = methicillin resistant *S. aureus*; CSE = carbapenem susceptible enterobacteriaceae; CRE = carbapenem resistant enterobacteriaceae; VSE = vancomycin susceptible enterococcus; VRE = vancomycin resistant enterococcus). Amplified bacterial genomic DNA using a one-pot pooled PCR is along the horizontal axis. Gene specific detection sets with crRNA guides are along the vertical axis. Stars indicate genes detected using whole genome sequencing. For more detailed strain information, see Table S3.

### Streamlined workflow for rapid testing and portable imaging

To demonstrate the potential for CARMEN to be broadly applied in the real world, we sought to address three key challenges. First, we wanted to reduce turnaround time and complexity of setup by eliminating the need for users to perform any detection droplet emulsion production, and to have a droplet-free sample loading approach to eliminate the need for benchtop droplet generation by the user. Second, we wanted to have a simple sample readout method that does not require intensive lab-based fluorescent microscopy.

To simplify the workflow, we preloaded and freeze-dried barcoded detection crRNAs in the microarray so that the only steps required to run an assay would be to load a single preamplified sample into the microarray and image the array, thus eliminating any user-performed, day-of-assay droplet/emulsion steps. To enable this, we employed microarrays with a circular well configuration. In a manufacturing step, microarray chips are loaded with barcoded droplets containing crRNAs corresponding to pathogens of interest, pre-imaged using fluorescent microscopy, and then freeze-dried using a lyophilizer. This pre-image thus enables crRNA identification based on well position and obviates the need for multiple color detection of barcodes in the field.

A droplet-free sample loading method was developed to introduce the pre-amplification reaction containing targets of interest into the freeze-dried, pre-loaded microarray (Figure 4b). After a period of incubation, sample readout is performed to determine which wells contained a crRNA guide that recognized a target from a corresponding pathogen, with the pre-imaged record reporting crRNA guide, and thus pathogen identity in each well. To enable simple sample readout, we employed a conventional cell phone camera for basic fluorescence imaging. The chip was illuminated using UV light and imaged after bringing the microchip wells into focus (no magnification needed.) The fluorescence image was recorded through a filter that blocked the UV excitation (see Methods.)

**Figure 4:**
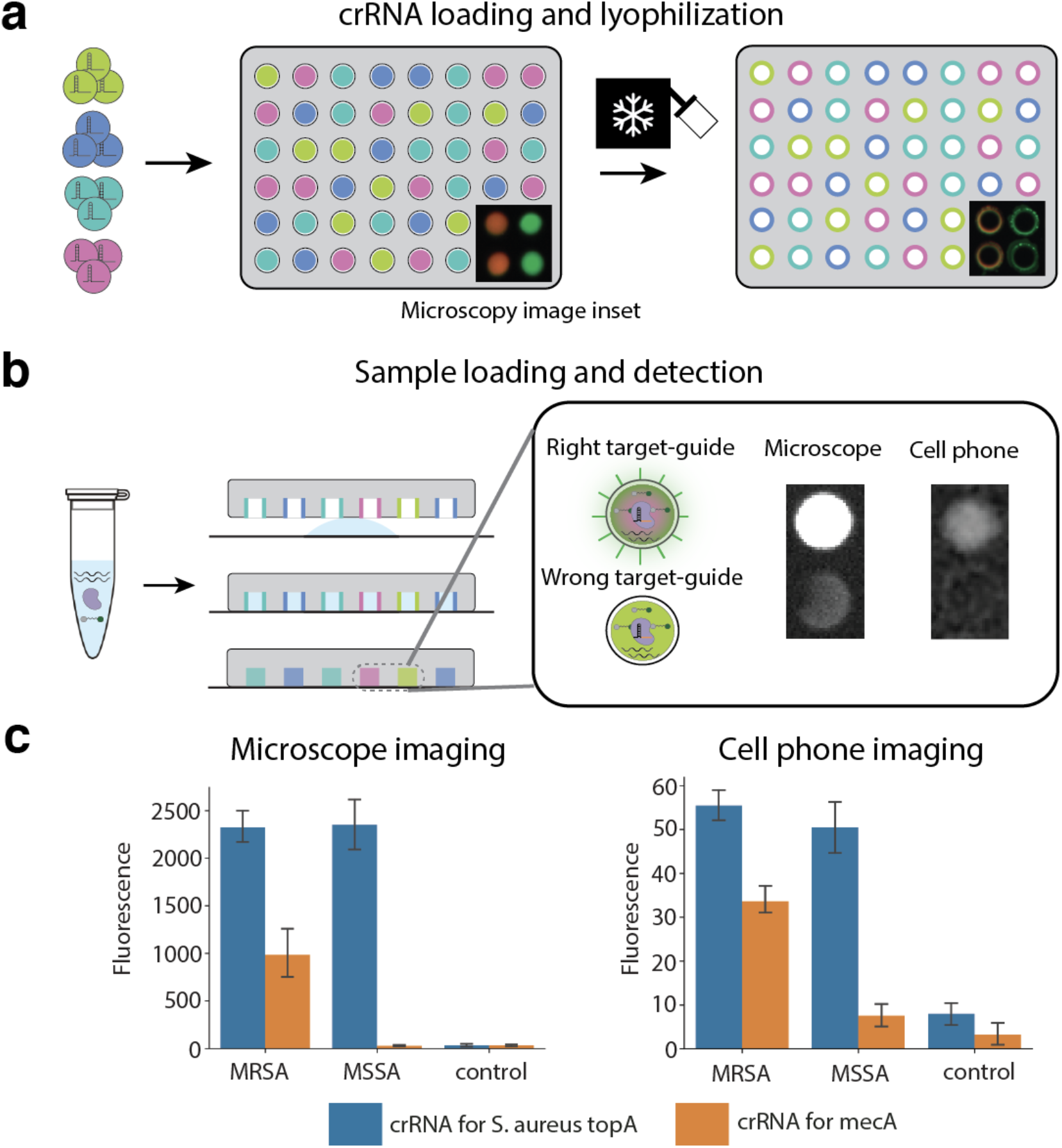
Droplet-free CARMEN v.2 for faster low-infrastructure testing. **a**, ‘Factory’ steps for pre-loaded array preparation and storage. A library of detection droplets containing crRNAs and fluorescent color codes are loaded into a droplet array with microwells engineered to capture one droplet each. The array is imaged to identify the crRNA present in each microwell and lyophilized to eliminate the emulsion’s oil and water components to stabilize the array for long-term storage. **b**, To run a droplet-free CARMEN v.2 test, the sample is processed and pre-amplified as usual, then mixed with the common SHERLOCK reagents and introduced as a homogeneous aqueous sample to a pre-loaded array. The sample fills the microwells in the array, which is sealed to a substrate. As the crRNA is solubilized, the detection reaction is initiated and positive wells are queried against the database of crRNA locations to provide the test readout. In this configuration, a monochromatic test imager is sufficient to read the reactivity in each well. **c**, Median fluorescence signal when imaging using a microscope and cell phone camera for preamplified MRSA and MSSA strains on a chip with crRNAs for the S. aureus topA and mecA genes (control = gDNA from unrelated bacterial species).

To demonstrate the performance of the entire workflow, we loaded a small microarray chip (~1000 wells) with crRNAs designed to detect the *S. aureus topA* gene as well as the methicillin-resistance encoding *mecA* gene (Figure 4a). We then loaded a pre-amplification reaction containing amplified genomic DNA from two different strains of *S. aureus* (MRSA and MSSA) or a control (*M. tuberculosis*), as well as Cas13 detection reagents (except crRNA), onto a chip pre-loaded with crRNAs for *S. aureus topA* and *mecA* genes. After incubation at 37°C for 3 hours, we imaged the chip using a cell phone camera. While the resulting signal-to-background ratio for the different target-crRNA combinations were lower than with conventional fluorescent microscopy imaging, the *S. aureus* crRNA nevertheless produced a significantly greater signal for the MSSA and MRSA strains compared with the control in this portable format. The *mecA* crRNA also showed a signal greater than background for the MRSA strain but not the MSSA strain (Figure 4c).

## Discussion

Molecular diagnostics are beginning to revolutionize infectious disease testing. PCR and isothermal amplification methods have been instrumental in our response to the ongoing COVID-19 pandemic. Despite these advances, culture remains the workhorse of bacterial diagnostics. One reason for this is that nucleic acid tests require a diagnostic hypothesis (eg. PCR) or can be complex and time-consuming (NGS). An idealized diagnostic would combine sensitivity, specificity, speed, and simplicity with comprehensive coverage across organisms and resistance determination^12^. Toward that end, we developed bCARMEN, which extends our previously published method, CARMEN^18^, to bacterial species identification and genotypic AST. One limitation of the earlier implementation of CARMEN^18^ for the viral panel was the need for multiple (up to 15), parallel pre-amplification reactions for each sample. Here we address the complexity imposed by multiple pre-amplification reactions by designing a one-pot universal preamplification step per sample that captures all desired targets, enabling simultaneous, multiplexed bacterial species and genotypic antibiotic resistance identification. We designed primers targeting conserved sequences in the *topA* gene, which enabled amplification of the *topA* gene segment across 52 bacterial species in a single amplification reaction per sample. We were then able to discriminate among species by designing crRNA guides against the 52 species-specific *topA* amplicons, taking advantage of the crRNA-dependent CRISPR-Cas13 binding specificity. Using the DropArray platform, we demonstrated the ability of bacterial CARMEN (bCARMEN) to uniquely identify 52 different species, including multiple strains of the same species. We similarly applied bCARMEN to detect a large number of genetic resistance determinants in a single detection step, and found it to be 100% accurate on a set of 27 clinical isolates. bCARMEN thus presents a novel way in which bacterial infections can be diagnosed.

In addition to diagnostic applications, our method demonstrates the potential for novel approaches to targeted sequencing and microbiome studies. Whereas we focused on a list of 52 pathogenic bacterial species, one can rapidly adapt our approach to detect a custom target set representing any defined set of bacteria, genes, or sequence variants of interest. Small scale panels for such applications have previously been reported^28^. Large target sets focused on gut, skin, or environmental microbes can all be straightforwardly designed and applied using CARMEN. Further, the specificity of CARMEN means that primer–guide sets can be designed for the desired level of taxonomic resolution, whether at the species or even strain level. These modular designs represent novel approaches to microbial genomics enabled by CARMEN.

Moving forward, we envision a strategy in which custom assay panels are designed based on clinical syndromes. For example, body site-specific panels focused on lower or upper respiratory pathogens, bloodstream pathogens, and urinary tract infections can be assembled to target pathogens known to cause disease at specific infection sites. The combinatorial nature of bCARMEN allows us to test many tens of patient samples at once, across a custom assay panel, thereby reducing reagent cost, while producing a distinct result for each patient-target combination. These panels would additionally combine pathogen identification with genotypic AST, thereby providing critical information for patient management.

Infrastructure requirements are another key challenge to current diagnostic tools for infectious disease. The version of CARMEN that we applied to viral identification significantly reduced turnaround time, requiring about seven hours, which is substantially faster than short-read NGS approaches. Key challenges to move CARMEN even further towards the point of care include: (1) reduction of setup time and complexity; (2) eliminating the need for benchtop droplet generation; (3) simple portable readout. Here, we exploited the robustness of Cas13 crRNAs to lyophilization to create a workflow involving preloaded chips to further reduce this time to less than three hours and eliminate the need for droplet generation. Further, we also demonstrated the ability to image the microwells using a widely available cell phone camera, thereby obviating the need for expensive fluorescent microscopes. By addressing these resource- and time-intensive steps in the workflow, we demonstrate the potential for CARMEN v.2 to be used in a point of care setting.

While we have demonstrated the ability to apply CRISPR-Cas13 and DropArray technologies to bacterial species identification, resistance gene detection, and addressed key challenges in point-of-care testing, there is still considerable work to be done to move from proof of principle to actual deployment in the real world. First, although our bacterial species identification panel demonstrated good specificity with an ability to discriminate each species, it was challenging to completely eliminate cross-talk in the assays for a few bacterial species in the two rounds of optimization we performed. Further improvement can be made by taking advantage of the modularity of our platform, for example to design multiple probes targeting different regions of the same amplicon in a way that they collectively offer additional specificity. The addition of more crRNA guides has minimal impact on overall assay operation, although interpretation of this more complicated readout may require additional computational support, integrated into analysis of the ensuing cell phone camera image. Second, while our resistance panel demonstrated the potential to detect resistance genes, genotypic resistance can also result from point mutations, depending on the organism and antibiotic. For example, pathogens predominantly acquire fluoroquinolone resistance through the acquisition of single nucleotide polymorphisms. While the specificity of Cas13 crRNAs allows for discrimination of such point mutations, this currently requires more sophisticated guide design and a more quantitative measurement of Cas13 activity for each of these guides^18^. Additionally, as the current assay detects genetic elements that confer resistance, limitations of genotypic AST also apply to our assay. Our current, incomplete knowledge of all genetic resistance elements limits the accuracy with which we can perform all AST; however, as our understanding of resistance mechanisms grows, so will our ability to predict resistance patterns based solely on genotype. Finally, although CARMEN v2 addresses key barriers to bringing CARMEN to the point-of-care, the challenge of rapid automated sample preparation from clinical material remains to be addressed.

Here we have demonstrated that bCARMEN both complements and advances our previously published CARMEN methodology applied to viruses. First, we detect a large panel of bacterial species by targeting a single locus, *topA*, in a one-pot amplification reaction to generate an amplicon enabling species-specific Cas13-based signal generation. We additionally demonstrate detection of a panel of clinically relevant resistance determinants. Finally, through the development of CARMEN v.2, we address key barriers towards bringing CARMEN to the point-of-care, including equipment and workflow complexity and time-to-result. It is our hope that through further development and testing, CARMEN will achieve its promise to transform clinical diagnostics and epidemiological surveillance of infectious diseases.

## Methods

### Primer and crRNA preparation

Individual primers were ordered from Integrated DNA Technologies and resuspended in nuclease-free water and stored at −20°C. crRNA guides were ordered as complementary ssDNA sequences with a T7 promoter binding sequence attached to the 5’-end. crRNA was synthesized in vitro using the HiScribe T7 High Yield RNA Synthesis Kit (New England Biolabs) by incubating the reaction at 37°C with T7 promoter primer (10μM) for 12 hours. In vitro transcribed product was then diluted down to a final concentration of 10 ng/μL of crRNA and quantified using a Nanodrop instrument (Thermo Scientific). crRNAs were stored at −80°C.

### Strain and gDNA preparation

A total of 113 strains across 52 species were obtained from local hospitals, collaborators, or strain collections (BEI, DSMZ, see Table S1). Strains included a combination of reference strains and clinical isolates. For strains obtained from collaborators or strain collections, strain identification was determined by the provider; for clinical isolates, this was performed using the standard workflow of CLIA certified, clinical microbiology laboratories. Where possible, gDNA from the strains was directly obtained. Remaining strains were cultured in liquid media to either mid-log or stationary phase, and DNA was extracted from liquid cultures using the DNeasy Blood and Tissue Kit (Qiagen.) Extracted gDNA was diluted down to 10^3^ genome equivalents per μL in nuclease-free water and used as input to amplification reactions.

### Synthetic targets

Synthetic targets were ordered from Integrated DNA Technologies and resuspended in nuclease-free water. Resuspended DNA was diluted to 10^3^ copies per μL and used as input to amplification reactions.

### Nucleic acid amplification

Amplification was performed by PCR using Q5 Hot Start polymerase (New England Biolabs) with total final primer concentration of 3.5 μM (individual primer concentrations varied depending on how many primers were pooled.) Unless otherwise stated, 30 cycles of PCR were performed using an annealing temperature of 65°C. Amplified samples were stored at −20°C until further use.

### Cas-13 detection reactions

Detection assays were performed with 45 nM purified *Leptotrichia wadei* Cas13a, 22.5 nM crRNA, 500 nM quenched fluorescent RNA reporter (RNAse Alert v2, Thermo Scientific), 2 μl murine RNase inhibitor (New England Biolabs) in nuclease assay buffer (40 mM Tris-HCl, 60 mM NaCl, pH 7.3) with 1 mM ATP, 1 mM GTP, 1 mM UTP, 1 mM CTP and 0.6 μl T7 polymerase mix (New England Biolabs)

### Barcoding, emulsification, and droplet pooling

Amplified samples were diluted 1:10 into nuclease-free water supplemented with 13.2 mM MgCl_2_ prior to barcoding with fluorescent dyes. Detection sets were prepared at 2.2x final concentration such that upon addition of barcoding dyes and merger with sample droplets, the final concentration of reagents would be 1x. Construction of fluorescent dye barcode sets has been described previously (citations). 2μL fluorescent dye barcode stocks were added to 18μL diluted sample mixture or detection mix for a final concentration of 2μM. Each amplified sample or detection mix received a distinct fluorescent barcode.

20 μL of each sample and detection mix were then emulsified into droplets using a BioRad QX200 droplet generator using fluorous oil (3M 7500, 70μL) containing 2% 008-fluorosurfactant (RAN Biotechnologies.)

For droplet pooling, a total emulsion volume of 150μL was used to load each standard chip; a total of μL was used to load each mChip. Half of this total volume was comprised of sample droplets, and the remaining half was of detection droplets. An equal volume of each sample and detection set added to each half total volume. The volume of each sample and detection mix varied on the experiment, but was typically between 5 and 12μL. The pooling step was rapid (<5min) and small molecule exchange does not alter color codes, as reported previously.

### Pooled droplet loading and imaging of microarrays

Loading and imaging of microarray chips was performed as described previously. Each chip was placed into an acrylic chip loader, suspended ~500μm the bottom acrylic surface, creating a between the chip and the loader. Fluorous oil (3M, 7500) was added to the flow space followed by the droplet pool. The loader was tilted above to move the droplet pool within the flow space until all the wells were filled. Fresh fluorous oil (3M, 7500) was added to wash off any excess droplets and the microarray chip was sealed using optically clear PCR film (MicroAmp, Applied Biosystems.)

Imaging of chips was done using fluorescent microscopy at 2x magnification (Nikon. MRD00025) and the following filter cubes: Alexa Fluor 555: Semrock SpGold-B; Alexa Fluor 594: Semrock 3FF03-575/25-25 + FF01-615/24-25; and Alexa Fluor 647: Semrock LF635-B. Pre-merge imaging was first performed to identify the contents of each well in the microarray. The droplets in each microwell were then merged by passing the tip of a corona treater (Model BD-20, Electro-Technic Products) over the PCR film. Merged droplets were imaged at 0, 1 hour, and 3 hour time points and incubated in a 37°C warm room in between imaging.

### Data analysis

Image data analysis was performed using custom Python scripts published previously. It consisted of the following parts: (1) pre-merge image analysis to identify the contents of each well in the microarray; (2) post-merge image analysis at each time to map pairs of droplets to microwells and measure reporter readout signal for that droplet pair. Heatmaps were then generated from the median fluorescence value of each crRNA-target pair.

### Primer and guide design for bacterial species panel

20 housekeeping genes from the 52 bacterial species were curated from multiple online databases (POGO-DB, NCBI, Ensembl.) Sequences from a total of 753 strains across 52 species (median = 10 strains per species) were used in the design process. For each of the 20 genes, 52 consensus sequences across strains from each species were determined and aligned using MUSCLE^29^. Nucleotide conservation was then plotted as a function of position. Conservation score at a given nucleotide position = occurrence of most prevalent nucleotide/52 (Figure S1.) One limitation of this method is that gaps in the alignment file produce low conservation scores. A low score therefore does not imply high diversity. topA, valS, and rpoB genes showed multiple regions with high degree of conservation flanking regions of variable conservation. Multiple pairs of degenerate primers were designed to target these conserved regions and their ability to amplify targets across species was evaluated using gel electrophoresis. Primers targeting a ~1000 b.p. region of the topA gene showed the most consistent amplification results across species, and this segment was chosen for further crRNA guide design to discriminate between species. Species specific primer pairs were ordered and pooled together (Table S2.)

For crRNA design, the target amplicon sequences for 753 strains across 52 species were aligned and fed into a diagnostic guide design algorithm called ADAPT. crRNA guides were designed to cover >95% of all strains within each target species, and be >3 SNPs apart from any region of non-target species. crRNAs were finally verified using a BLAST search to ensure no cross-reactivity against the human genome (>5 SNPs apart), as well as other bacterial organisms (>3 SNPs apart.)

### Primer and guide design for bacterial resistance panel

For each resistance gene, multiple gene sequences were obtained from the Comprehensive Antibiotic Resistance Database (median = 7 sequences per resistance gene.) Gene sequences were then aligned and primers and crRNA guides were visually designed based on the design strategy. A blast search was performed on the primers and guides to ensure no cross-reactivity.

### Pre-test setup for droplet-free assay

The “factory” step of the droplet-free assay involved preparation of dilution of crRNA guides in nuclease-free water to a final concentration of 225 nM. Barcoding, emulsification, and pooling was then performed as before for just the crRNA droplets. Pooled droplets were then loaded on a microarray chip as before, but using a microarray chip with a different well configuration. Each well was circular and large enough to hold a single droplet. The droplet was loaded and sealed as before, imaged using fluorescent microscopy to determine the identity of each microwell, and immediately placed in a −80°C freezer for at least two hours. After this, the frozen chip was transferred onto a lyophilizer (ThermoModulyo) overnight. The freeze-dried chip was then stored at room temperature for up to 24 hours before being used.

### Target preparation, loading, and imaging of microarrays

Amplified target was diluted 1:20 with 9 mM MgCl2, 45 nM purified *Leptotrichia wadei* Cas13a, 500 nM quenched fluorescent RNA reporter (RNAse Alert v2, Thermo Scientific), 2 μl murine RNase inhibitor (New England Biolabs) in nuclease assay buffer (40 mM Tris-HCl, 60 mM NaCl, pH 7.3) with 1 mM ATP, 1 mM GTP, 1 mM UTP, 1 mM CTP and 0.6 μl T7 polymerase mix (New England Biolabs.) This target mix was then loaded on to the chip by placing 10 μL of the mix on the sticky side of the PCR film (MicroAmp, Applied Biosystems) and pressing down the freeze-dried microarray chip on this mixture (Figure 4B, S6A.) A weight was placed on the chip until the PCR film was stuck and the chip was incubated at 37°C for 3 hours.

After incubation, the reporter channel of the chip was imaged using a standard gel illuminator (E-gel Power Snap, Invitrogen) and a cellphone camera using the “macro” focus option (Figure S6B.) For cell phones without this option, a small lens band (Easy-Macro) was used to achieve focu of the microwells. The captured image was correlated with the pre-test image and reporter signal for each crRNA-target pair was determined manually using ImageJ.

### Ethical approval and informed consent

Discarded clinical samples from Massachusetts General Hospital and Brigham and Women’s Hospital were obtained under waiver of consent due to exclusive focus on pathogen and not host contents, as approved by the Partners HealthCare Institutional Review Board that governs both institutions, under protocol number 2015P002215.

