## Supplemental Figures and Tables for "Multiplexed detection of bacterial nucleic acids using Cas13 in droplet microarrays"

**Supplementary figures and tables:**

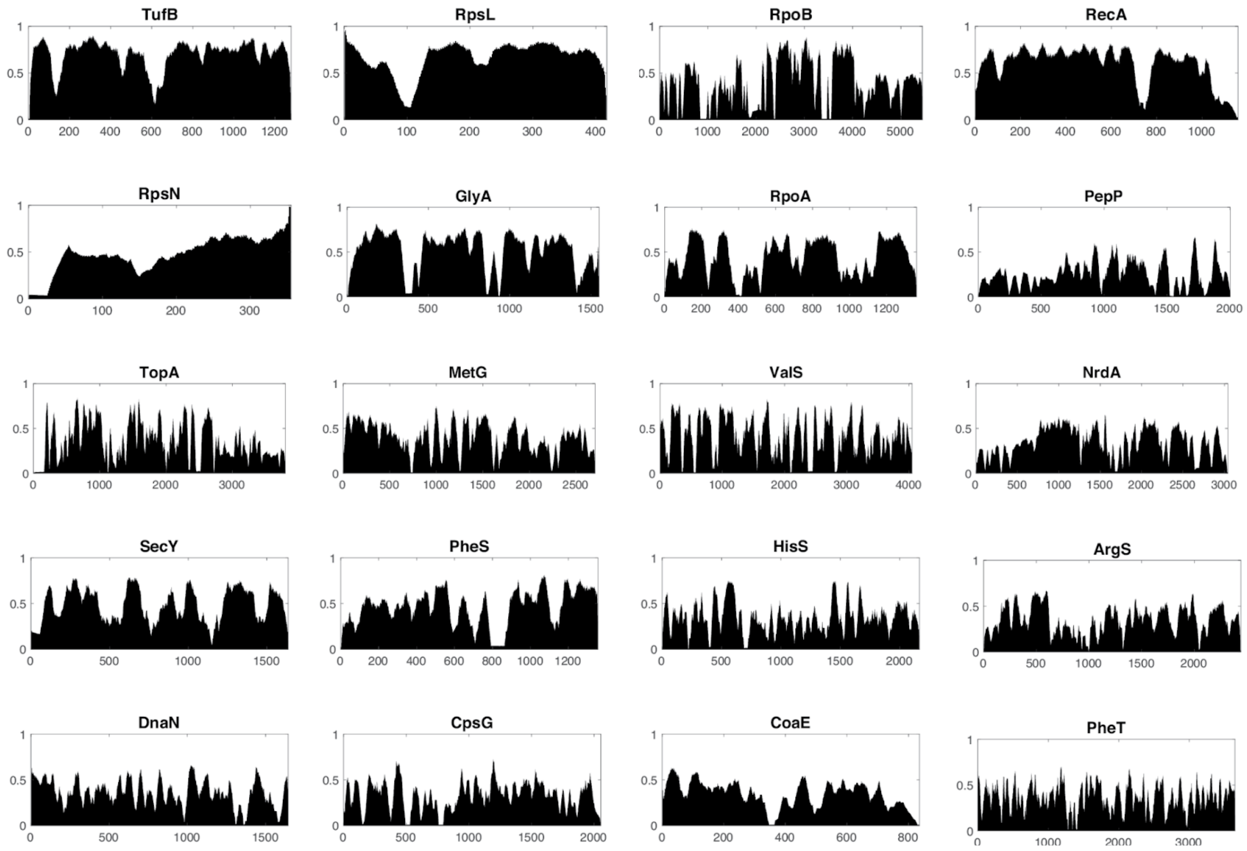


**Figure S1: Nucleotide conservation of housekeeping genes across bacterial species**
Conservation as a function nucleotide position across 52 bacterial species plotted for 20 housekeeping genes. Conservation score at a given nucleotide position = occurrence of most prevalent nucleotide / 52.

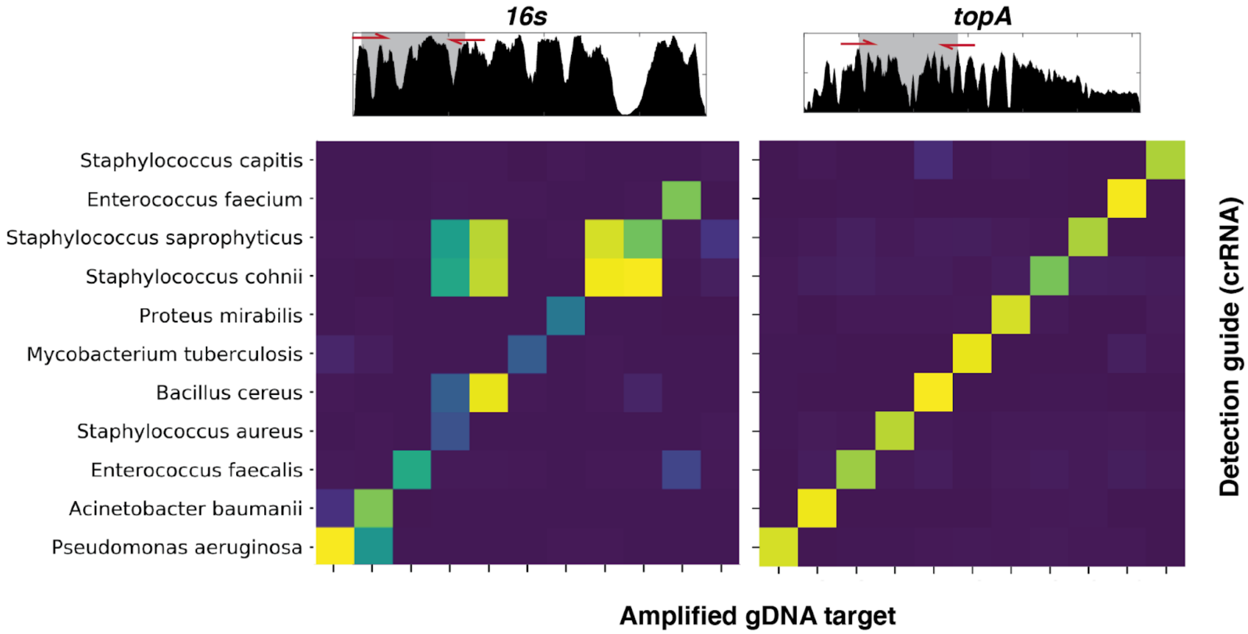


**Figure S2: Comparison of performance of 16s vs. topA crRNA guides** Heatmap showing fluorescent signal from crRNAs targeting a subset of species at 16s vs. topA loci. Genomic loci amplified by primers is depicted above the heatmap.

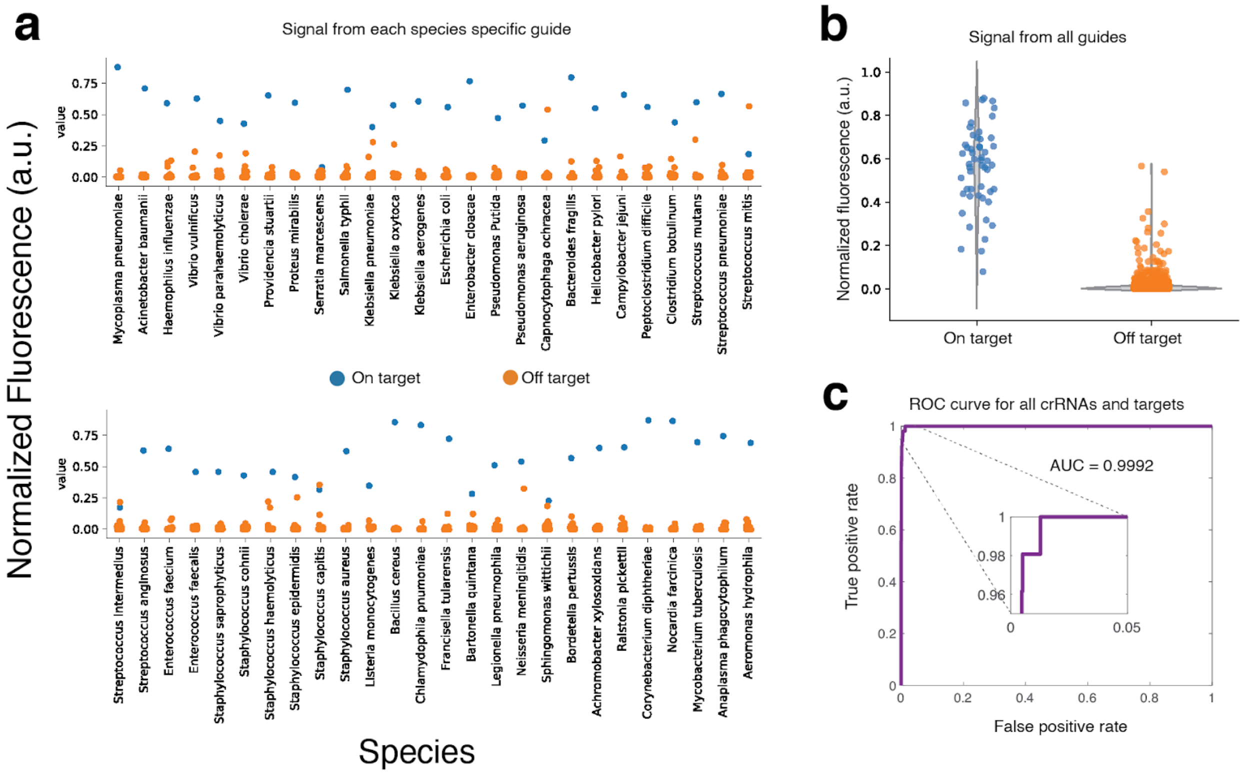


**Figure S3: crRNA guide performance in the bacterial panel a**, Fluorescent signal for each species-specific crRNA plotted for the target species (blue, on target) and all non-target species (orange, off target.) **b**, Fluorescent signal from all crRNA guides plotted for target species (blue, on target) and all non-target species (orange, off target.) **c**, Receiver operator characteristics curve plotted for on-target vs. off-target signals from all crRNA guides (AUC = 0.9992)

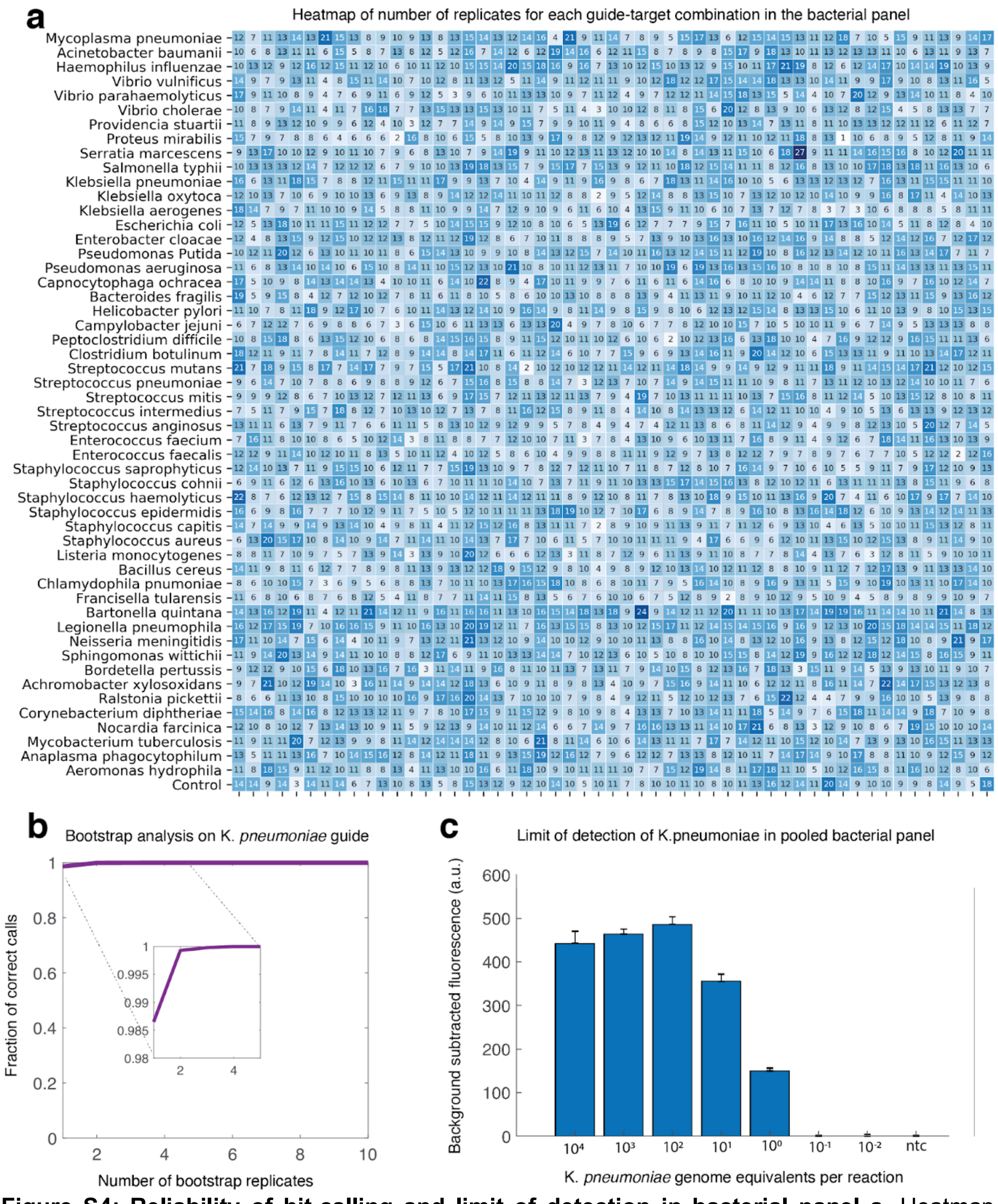


**Figure S4: Reliability of hit-calling and limit of detection in bacterial panel a**, Heatmap depicting the number of replicates for each crRNA-target pair in the bacterial panel data (Figure 2d) Median = 10.8 **b**, Bootstrap analysis on K. *pneumoniae* guide showing call confidence (fraction of correct calls) as a function of number of replicates. **c**, Limit of detection of the bacterial panel assay

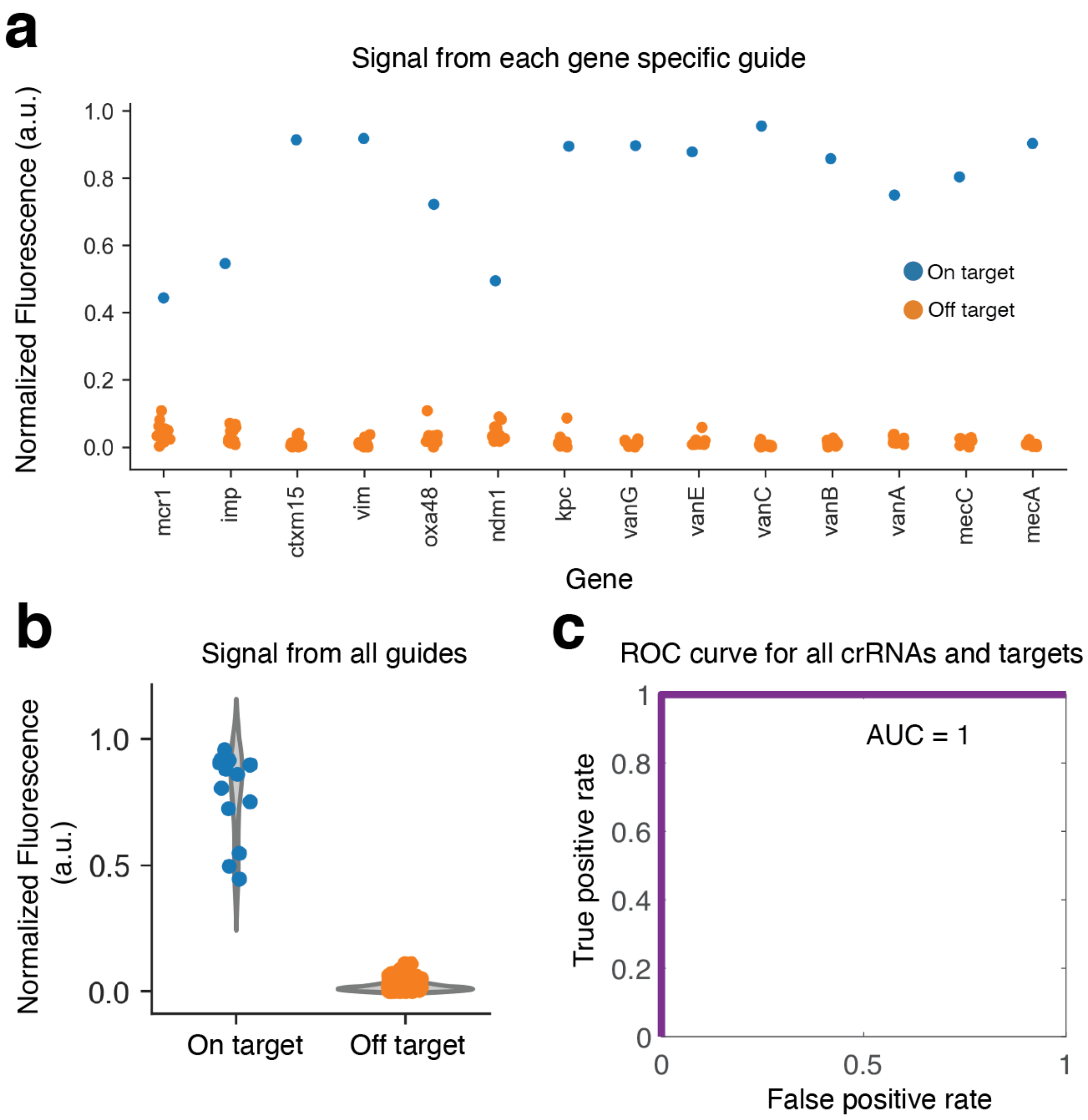


**Figure S5: crRNA guide performance in the resistance panel a**, Fluorescent signal for each species-specific crRNA plotted for the target gene (blue, on target) and all non-target genes (orange, off target.) **b**, Fluorescent signal from all crRNA guides plotted for target gene (blue, on target) and all non-target genes (orange, off target.) **c**, Receiver operator characteristics curve plotted for on-target vs. off-target signals from all crRNA guides (AUC = 1)

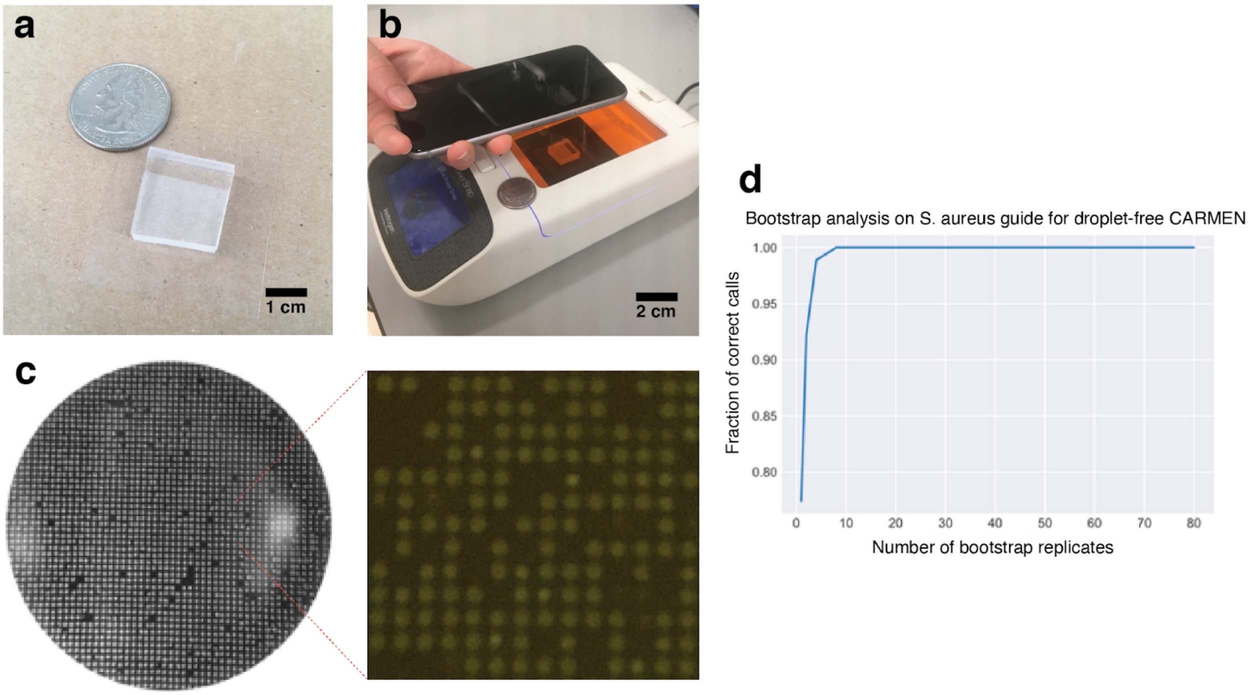


**Figure S6: Droplet-free assay setup and bootstrap analysis a**, Freeze-dried chip loaded with target mix on PCR film with an American quarter dollar coin shown for scale. **b**, Imaging setup used for droplet-free assay. **c**, Cell phone image of microwells with a zoomed in image showing the droplets close up. d, Bootstrap analysis on S. aureus topA guide plotting the fraction of correct calls as a function of number of replicates.

**Table S1: Strain information for bacterial species panel**

| Species | Tested strain name | Source | Number of strains used in guide design |
|---|---|---|---|
| Mycoplasma pneumoniae | DSM 23978 | DSMZ | 4 |
| Acinetobacter baumanii | RB197 | BWH clinical isolate | 14 |
| Haemophilus influenzae | DSM4690 | DSMZ | 20 |
| Vibrio vulnificus | DSM10143 | DSMZ | 5 |
| Vibrio parahaemolyticus | RIMD 2210633 | Goldberg lab (MGH) | 12 |
| Vibrio cholerae | | MGH clinical isolate | 98 |
| Providencia stuartii | DSM4539 | DSMZ | 2 |
| Proteus mirabilis | RB037 | BWH clinical isolate | 5 |
| Serratia marcescens | BWH_23 | BWH clinical isolate | 10 |
| Salmonella typhii | NR-514 | BEI | 45 |
| Klebsiella pneumoniae | RB011 | BWH clinical isolate | 15 |
| Klebsiella oxytoca | RB078 | BWH clinical isolate | 10 |
| Klebsiella aerogenes | RB251 | MGH clinical isolate | 3 |
| Escherichia coli | RB001 | MGH clinical isolate | 53 |
| Enterobacter cloacae | RB250 | MGH clinical isolate | 15 |
| Pseudomonas putida | WCS358 | Lory lab | 10 |
| Pseudomonas aeruginosa | RB019 | BWH clinical isolate | 16 |
| Capnocytophaga ochracea | DSM7271 | DSMZ | 2 |
| Bacteroides fragilis | DSM2151 | DSMZ | 3 |
| Helicobacter pylori | CPY6081 | BEI | 51 |
| Campylobacter jejuni | NR-3057 | BEI | 14 |
| Peptoclostridium difficile | CD104 | BEI | 4 |
| Clostridium botulinum | NR-2713 | BEI | 13 |
| Streptococcus mutans | DSM20523 | DSMZ | 10 |
| Streptococcus pneumoniae | D39 | Lipsitch lab | 24 |
| Streptococcus mitis | Spar10 | BEI | 27 |
| Streptococcus intermedius | F0413 | BEI | 3 |
| Streptococcus anginosus | F0211 | BEI | 8 |
| Enterococcus faecium | RB029 | BWH clinical isolate | 4 |
| Enterococcus faecalis | RB027 | BWH clinical isolate | 5 |
| Staphylococcus saprophyticus | RB497 | Wadsworth lab clinical isolate | 2 |
| Staphylococcus cohnii | RB493 | Wadsworth lab clinical isolate | 3 |
| Staphylococcus haemolyticus | DNF00585 | BEI | 3 |
| Staphylococcus epidermidis | RB494 | Wadsworth lab clinical isolate | 8 |

| | | | |
|---|---|---|---|
| Staphylococcus capitis | RB492 | Wadsworth lab clinical isolate | 3 |
| Staphylococcus aureus | Newman | BEI | 40 |
| Listeria monocytogenes | 10403s | Goldberg lab (MGH) | 29 |
| Bacillus cereus | TOR16585 | BEI | 37 |
| Chlamydophila pnumoniae | DSM19748 | DSMZ | 5 |
| Francisella tularensis | NR-3015 | BEI | 10 |
| Bartonella quintana | JK39 | BEI | 3 |
| Legionella pneumophila | Lp01 JK40 | Isberg lab (Tufts) | 29 |
| Neisseria meningitidis | NM3222 | BEI | 12 |
| Sphingomonas wittichii | DSM6014 | DSMZ | 1 |
| Bordetella pertussis | H921 | BEI | 3 |
| Achromobacter xylosoxidans | DSM2042 | DSMZ | 2 |
| Ralstonia pickettii | DSM6297 | DSMZ | 3 |
| Corynebacterium diphtheriae | DSM44123 | DSMZ | 11 |
| Nocardia farcinica | DSM43665 | DSMZ | 1 |
| Mycobacterium tuberculosis | H37Rv | Lab | 21 |
| Anaplasma phagocytophilum | NR-51150 | BEI | 20 |
| Aeromonas hydrophila | DSM30187 | DSMZ | 2 |

**Table S2: Primer and guide sequences for bacterial species panel**

| Species | Forward Primer Sequence | Reverse Primer Sequence | Guide (reverse complement with T7 promoter attached) |
|---|---|---|---|
| Mycoplasma pneumoniae | gaaatTAATACGACTCACTATAGGGAGACCGTGAGGGAGAGGCAATTTCCTGACA | AGTTGGGTGAATTGCTTCATGGGCGTCTTG | AAGGACAAGTCGAAGTGCCAACGCATTAGTTTTAGTCCCCTTCGTTTTTGGGGTAGTCTAAATCCCCTATAGTGAGTCGTATTAatttc |
| Acinetobacter baumanii | gaaatTAATACGACTCACTATAGGGGGATAGAGAAGGGGAAGCAATCGCTTGGCA | TTGATGGACGAATAGCTTCGTGGGCTTCTTG | CAGCCAACACGTCTCGACTTAAACCGTGGTTTTAGTCCCCTTCGTTTTTGGGGTAGTCTAAATCCCCTATAGTGAGTCGTATTAatttc |
| Haemophilus influenzae | gaaatTAATACGACTCACTATAGGGGATAGAGAAGGGGAAGCGATTGCGTGGCA | TGATGGACGAATAGCCTCATGAGCCTCTTG | ACGATGATCGTTTTAGTCGCGTGGTGTTGTTTTAGTCCCCTTCGTTTTTGGGGTAGTCTAAATCCCCTATAGTGAGTCGTATTAatttc |
| Vibrio vulnificus | gaaatTAATACGACTCACTATAGGGACCGCGAGGGGGAAGCTATTGCGTGGCA | TGAAGGACGAATCGCTTCGTGTGCTTCTTG | AGCTATTGCGTGGCACCTTCGTGAGATCGTTTTAGTCCCCTTCGTTTTTGGGGTAGTCTAAATCCCCTATAGTGAGTCGTATTAatttc |
| Vibrio parahaemolyticus | gaaatTAATACGACTCACTATAGGGACCGCGAGGGGGAAGCAATCGCATGGCA | AGATGGACGAATCGCTTCGTGCGCTTCTTG | CGATGAAGAGCGATACAAACGAGTAGTCGTTTTAGTCCCCTTCGTTTTTGGGGTAGTCTAAATCCCCTATAGTGAGTCGTATTAatttc |
| Vibrio cholerae | gaaatTAATACGACTCACTATAGGGACCGCGAGGGGGAAGCAATCGCATGGCA | AGATGGACGAATCGCTTCGTGCGCTTCTTG | CGATGAAAAGCGATATAAACGAGTGGTAGTTTTAGTCCCCTTCGTTTTTGGGGTAGTCTAAATCCCCTATAGTGAGTCGTATTAatttc |
| Providencia stuartii | gaaatTAATACGACTCACTATAGGGGACCGCGAAGGGGAAGCTATTGCATGGCA | GAAGGACGGATAGCTTCGTGCGCTTCTTG | CATGGATCGCGTAGTAGGTTATATGGTAGTTTTAGTCCCCTTCGTTTTTGGGGTAGTCTAAATCCCCTATAGTGAGTCGTATTAatttc |
| Proteus mirabilis | gaaatTAATACGACTCACTATAGGGACCGCGAAGGAGAAGCTATCGCGTGGCA | AGAAGGACGAATAGCTTCGTGCGCTTCTTG | ATGCCATTACACAAGCATTCCAAACACCGTTTTAGTCCCCTTCGTTTTTGGGGTAGTCTAAATCCCCTATAGTGAGTCGTATTAatttc |
| Serratia marcescens | gaaatTAATACGACTCACTATAGGGACCGCGAAGGGGAAGCCATTGCCTGGCA | AAGGACGAATCGCCTCGTGCGCTTCCTG | CGATCTGCTGGCGAAGGGTGAAACCGCGGTTTTAGTCCCCTTCGTTTTTGGGGTAGTCTAAATCCCCTATAGTGAGTCGTATTAatttc |
| Salmonella typhii | gaaatTAATACGACTCACTATAGGGACCGCGAAGGGGAAGCCATTGCATGGCA | AAGGGCGGATCGCTTCGTGCGCTTCCTG | TTGTCGGCAGGTCGCGTTCAGTCTGTCGGTTTTAGTCCCCTTCGTTTTTGGGGTAGTCTAAATCCCCTATAGTGAGTCGTATTAatttc |
| Klebsiella pneumoniae | gaaatTAATACGACTCACTATAGGGACCGCGAAGGGGAAGCCATTGCCTGGCA | GAAGGACGAATCGCTTCGTGCGCTTCCTG | CAGCCGTGATGAGACCATGGCGGCGGTAGTTTTAGTCCCCTTCGTTTTTGGGGTAGTCTAAATCCCCTATAGTGAGTCGTATTAatttc |
| Klebsiella oxytoca | gaaatTAATACGACTCACTATAGGGACCGCGAAGGGGAAGCCATCGCATGGCA | ATGGACGAATCGCCTCGTGCGCTTCCTG | GTCACGCATAACGGCGATAAGCCGTTCCGTTTTAGTCCCCTTCGTTTTTGGGGTAGTCTAAATCCCCTATAGTGAGTCGTATTAatttc |
| Klebsiella aerogenes | gaaatTAATACGACTCACTATAGGGACCGCGAAGGGGAAGCCATTGCCTGGCA | GAAGGACGAATCGCTTCGTGCGCTTCCTG | TATCGGTGGTGATGAGCAACGGTACAGCGTTTTAGTCCCCTTCGTTTTTGGGGTAGTCTAAATCCCCTATAGTGAGTCGTATTAatttc |
| Escherichia coli | gaaatTAATACGACTCACTATAGGGACCGCGAAGGGGAAGCCATTGCATGGCA | GAAGGACGAATCGCTTCGTGCGCTTCCTG | ATTGGTGGTGATGATGCGCGCTATAGCCGTTTTAGTCCCCTTCGTTTTTGGGGTAGTCTAAATCCCCTATAGTGAGTCGTATTAatttc |
| Enterobacter cloacae | gaaatTAATACGACTCACTATAGGGACCGCGAAGGGGAAGCCATTGCCTGGCA | GAAGGACGAATGGCTTCGTGCGCTTCCTG | AAAGAATGCGATTCGTCAGGCGTTTGAGGTTTTAGTCCCCTTCGTTTTTGGGGTAGTCTAAATCCCCTATAGTGAGTCGTATTAatttc |
| Pseudomonas putida | gaaatTAATACGACTCACTATAGGGACCGCGAAGGGGAAGCCATTGCCTGGCA | AAGGCCGAATCGCTTCGTGCGCCTCCTG | GGCACCGCCAAGAACGCCAAGGTGCGTTGTTTTAGTCCCCTTCGTTTTTGGGGTAGTCTAAATCCCCTATAGTGAGTCGTATTAatttc |
| Pseudomonas aeruginosa | gaaatTAATACGACTCACTATAGGGCGCGAAGGGGAGGCCATCGCCTGGCA | AGGACGGATCGCCTCGTGCGCTTCCTG | TTCTCCCAGCCCGGCGAGCTCGATATCAGTTTTAGTCCCCTTCGTTTTTGGGGTAGTCTAAATCCCCTATAGTGAGTCGTATTAatttc |
| Capnocytophaga ochracea | gaaatTAATACGACTCACTATAGGGAAGACCGTGAAGGAGAAGCTATTGCGTTTCA | GTAGGACGAATGGCTTCGTGCGCCTCTTG | ACTCTCAGCAGGACGCGTTCAGTCGGTAGTTTTAGTCCCCTTCGTTTTTGGGGTAGTCTAAATCCCCTATAGTGAGTCGTATTAatttc |
| Bacteroides fragilis | gaaatTAATACGACTCACTATAGGGGACCGGGAGGGAGAAGCCATTTCATGGCA | TCGGGCGGATAGCTTCGTGCGCTTCCTG | GCAGGCACGCCGTATCTTAGACCGTATCGTTTTAGTCCCCTTCGTTTTTGGGGTAGTCTAAATCCCCTATAGTGAGTCGTATTAatttc |
| Helicobacter pylori | gaaatTAATACGACTCACTATAGGGAGACAGAGAGGGGGAAGCGATAGGCTATCA | GTGGGCCTGATCGCTTCATGGGCTTCTTG | TTTCATGAGATCACGCAAAATGCGATTTGTTTTAGTCCCCTTCGTTTTTGGGGTAGTCTAAATCCCCTATAGTGAGTCGTATTAatttc |
| Campylobacter jejuni | gaaatTAATACGACTCACTATAGGGAGGATAGAGAAGGAGAGGCTATAGCCTATCA | TTGTAGGGCGTATAGCTTCATGGGCTTCTTG | GCGTGTTCAAAGTGCGGCTTTAAAGATCGTTTTAGTCCCCTTCGTTTTTGGGGTAGTCTAAATCCCCTATAGTGAGTCGTATTAatttc |
| Peptoclostridium difficile | gaaatTAATACGACTCACTATAGGGCCTGATAGAGAAGGTGAAGCTATATCATGGCA | AGGTTGGTCTTATTGCCTCATGGGCATCTTG | ACAGGCAAGAAGAGTGCTAGACAGACTTGTTTTAGTCCCCTTCGTTTTTGGGGTAGTCTAAATCCCCTATAGTGAGTCGTATTAatttc |
| Clostridium botulinum | gaaatTAATACGACTCACTATAGGGCGGATAGAGAGGGAGAAGCTATTTCTTGGCA | TGGTTGGTCTTATAGCTTCATGGGCATCCTG | AGATTAGTTGGATATAAAATAAGCCCTAGTTTTAGTCCCCTTCGTTTTTGGGGTAGTCTAAATCCCCTATAGTGAGTCGTATTAatttc |
| Streptococcus mutans | gaaatTAATACGACTCACTATAGGGGGACCGTGAAGGAGAAGCGATTTCTTGGCA | AGACGGACGAATAGCCTCATGAGCATCCTG | CGTAATGCACCGCTTCCTTATACGACATGTTTTAGTCCCCTTCGTTTTTGGGGTAGTCTAAATCCCCTATAGTGAGTCGTATTAatttc |
| Streptococcus pneumoniae | gaaatTAATACGACTCACTATAGGGGGACCGTGAAGGAGAAGCGATTTCTTGGCA | GACGGACGAATAGCCTCATGGGCATCCTG | GTTCAGTCCATTGCCCTTAAACTCATCAGTTTTAGTCCCCTTCGTTTTTGGGGTAGTCTAAATCCCCTATAGTGAGTCGTATTAatttc |
| Streptococcus mitis | gaaatTAATACGACTCACTATAGGGGGACCGTGAAGGAGAAGCGATTTCTTGGCA | AGACGGACGAATAGCCTCATGAGCATCCTG | CGCGTTCAGTCTATTGCGCTTAAGTTAAGTTTTAGTCCCCTTCGTTTTTGGGGTAGTCTAAATCCCCTATAGTGAGTCGTATTAatttc |
| Streptococcus intermedius | gaaatTAATACGACTCACTATAGGGCAGATCGAGAAGGAGAAGCTATTTCTTGGCA | TCGATGGACGAATCGCTTCATGAGCATCTTG | AGAAGGGGTTATCGGCTGGACGTGTACAGTTTTAGTCCCCTTCGTTTTTGGGGTAGTCTAAATCCCCTATAGTGAGTCGTATTAatttc |
| Streptococcus anginosus | gaaatTAATACGACTCACTATAGGGGGACCGTGAAGGAGAAGCTATTTCTTGGCA | GACGGACGAATCGCTTCATGGGCGTCTTG | TAAAGAACCGCGCCAGATTGACATGGATGTTTTAGTCCCCTTCGTTTTTGGGGTAGTCTAAATCCCCTATAGTGAGTCGTATTAatttc |
| Enterococcus faecium | gaaatTAATACGACTCACTATAGGGGGATAGAGAAGGAGAAGCGATTGCTTGGCA | GTTGGACGCACAGCTTCATGGGCATCTTG | AGTGCAGGACGTGTCCAATCTGTCGCTTGTTTTAGTCCCCTTCGTTTTTGGGGTAGTCTAAATCCCCTATAGTGAGTCGTATTAatttc |
| Enterococcus faecalis | gaaatTAATACGACTCACTATAGGGGGACCGAGAAGGTGAAGCAATTGCTTGGCA | GACGGACGAACCGCTTCATGGGCATCTTG | CGTTTAGTTGGATACTCGATTAGTCCTAGTTTTAGTCCCCTTCGTTTTTGGGGTAGTCTAAATCCCCTATAGTGAGTCGTATTAatttc |
| Staphylococcus saprophyticus | gaaatTAATACGACTCACTATAGGGCGACCGTGAAGGTGAAGCTATTGCTTGGCA | AGACGGTCTGATGGCTTCATGGGCATCTTG | ACAAGCACGACGTATTCTAGATCGTTTAGTTTTAGTCCCCTTCGTTTTTGGGGTAGTCTAAATCCCCTATAGTGAGTCGTATTAatttc |
| Staphylococcus cohnii | gaaatTAATACGACTCACTATAGGGCGATCGTGAAGGTGAGGCAATTGCTTGGCA | TAGATGGTCTAATAGCTTCGTGGGCATCTTG | GTGGATGCACAACAGGCTAGAAGAATTTGTTTTAGTCCCCTTCGTTTTTGGGGTAGTCTAAATCCCCTATAGTGAGTCGTATTAatttc |
| Staphylococcus haemolyticus | gaaatTAATACGACTCACTATAGGGCGACCGTGAAGGTGAAGCAATTGCTTGGCA | TTGAAGGTCTAATAGCCTCATGGGCATCTTG | GCTGTCTGCAGGGCGAGTTCAATCTGTAGTTTTAGTCCCCTTCGTTTTTGGGGTAGTCTAAATCCCCTATAGTGAGTCGTATTAatttc |
| Staphylococcus epidermidis | gaaatTAATACGACTCACTATAGGGTGACCGTGAAGGTGAAGCGATTGCTTGGCA | CTAGTAGGTCTAATAGCTTCGTGAGCATCTTG | GCTGGGAGAGTTCAGTCAGTTGCTTTACGTTTTAGTCCCCTTCGTTTTTGGGGTAGTCTAAATCCCCTATAGTGAGTCGTATTAatttc |
| Staphylococcus capitis | gaaatTAATACGACTCACTATAGGGGACCGTGAAGGTGAAGCGATTGCCTGGCA | TTGTAGGTCTAATCGCTTCATGGGCATCTTG | GTTCAGTCAGTAGCTCTTCGTTTAGTTAGTTTTAGTCCCCTTCGTTTTTGGGGTAGTCTAAATCCCCTATAGTGAGTCGTATTAatttc |
| Staphylococcus aureus | gaaatTAATACGACTCACTATAGGGCGACCGTGAAGGTGAAGCAATTGCTTGGCA | TTGAAGGTCTAATAGCCTCATGGGCATCTTG | AGAGCTTGAAGATTCTAAAGAAAATCGCGTTTTAGTCCCCTTCGTTTTTGGGGTAGTCTAAATCCCCTATAGTGAGTCGTATTAatttc |
| Listeria monocytogenes | gaaatTAATACGACTCACTATAGGGAGACCGCGAAGGAGAAGCAATTGCATGGCA | TTGTAGGTCTGATTGCTTCATGGGCATCTTG | TAGAGTTAGATCAATCAGACAAACTACGGTTTTAGTCCCCTTCGTTTTTGGGGTAGTCTAAATCCCCTATAGTGAGTCGTATTAatttc |
| Bacillus cereus | gaaatTAATACGACTCACTATAGGGAGACCGCGAAGGAGAAGCTATTGCTTGGCA | AAGTAGGACGAATTGCCTCATGCGCATCTTG | CAAGCAAGACGTATACTAGATCGTCTTGGTTTTAGTCCCCTTCGTTTTTGGGGTAGTCTAAATCCCCTATAGTGAGTCGTATTAatttc |
| Chlamydophila pnumoniae | gaaatTAATACGACTCACTATAGGGTGATAGAGAAGGAGAGGCAATTGCCTGGCA | AGTGGGACGTATGGCTTCGTGAGCATCTTG | CTGGCACATCGCGAATCAGCTTCCTGACGTTTTAGTCCCCTTCGTTTTTGGGGTAGTCTAAATCCCCTATAGTGAGTCGTATTAatttc |
| Francisella tularensis | gaaatTAATACGACTCACTATAGGGCCAGATAGAGAAGGTGAAGCTATATCATGGCA | TGTGACACGGATAGCTTCGTGTGCTTCTTG | ATCTATCGTGTCACATTTAACGAAATCAGTTTTAGTCCCCTTCGTTTTTGGGGTAGTCTAAATCCCCTATAGTGAGTCGTATTAatttc |
| Bartonella quintana | gaaatTAATACGACTCACTATAGGGCTGATCGTGAAGGGGAAGCTATTTCATGGCA | TTGGCCGGATGGCTTCATGCGCTTCCTG | TATCGATACATCACTTGTAGATGCCTACGTTTTAGTCCCCTTCGTTTTTGGGGTAGTCTAAATCCCCTATAGTGAGTCGTATTAatttc |

| | | | |
|---|---|---|---|
| Legionella pneumophila | gaaatTAATACGACTCACTATAGGGTGATAGAGAGGGAGAAGCTATCTCGTGGCA | AAGTTGGCCTAATGGCTTCATGTGCTTCCTG | ATCGGCCGGACGAGTACAAAGCCCTGCCGTTTTAGTCCCCTTCGTTTTTGGGGTAGTCTAAATCCCCTATAGTGAGTCGTATTAatttc |
| Neisseria meningitidis | gaaatTAATACGACTCACTATAGGGGATAGGGAAGGCGAAGCCATTTCCTGGCA | TCGGACGGATGGCTTCGTGCGCTTCTTG | AGGATGCTGTGCGCAAACTCGGCTTCACGTTTTAGTCCCCTTCGTTTTTGGGGTAGTCTAAATCCCCTATAGTGAGTCGTATTAatttc |
| Sphingomonas wittichii | gaaatTAATACGACTCACTATAGGGCGCGAGGGGGAGGCGATCAGCTGGCA | CGGGCGGATCGCCTCATGCGCTTCCTG | TCACCTTCAACGCGATCACCAAGGCCGCGTTTTAGTCCCCTTCGTTTTTGGGGTAGTCTAAATCCCCTATAGTGAGTCGTATTAatttc |
| Bordetella pertussis | gaaatTAATACGACTCACTATAGGGGGCGCGAAGGCGAACTGATCTTCCGCTA | TGTTGTCGAAGATGCGCCGGTTGGGCTTG | GCAGACGCCCACGCTGGCCATCGTCAATGTTTTAGTCCCCTTCGTTTTTGGGGTAGTCTAAATCCCCTATAGTGAGTCGTATTAatttc |
| Achromobacter xylosoxidans | gaaatTAATACGACTCACTATAGGGGGCGCGAGGGCGAACTGATCTTCCGCTA | TGTTGTCGAAGATGCGGCGGTTCGGCTTG | AGCGCGAGGAACGCATCCGCCGCTTCGTGTTTTAGTCCCCTTCGTTTTTGGGGTAGTCTAAATCCCCTATAGTGAGTCGTATTAatttc |
| Ralstonia pickettii | gaaatTAATACGACTCACTATAGGGGGCGCGAAGGCGAGTTGATCTTCCGCCT | CTGTTGTCGAAGATCCGCTTGTTCGGCTTG | GATCGTGGTGGAGCGCGAAGAGAAGATCGTTTTAGTCCCCTTCGTTTTTGGGGTAGTCTAAATCCCCTATAGTGAGTCGTATTAatttc |
| Corynebacterium diphtheriae | gaaatTAATACGACTCACTATAGGGACCGCGAGGGTGAAGCCATCGCTTGGCA | CTGGACGGATTGCCTCGTGGGCTTCTTG | CTACGGCTACGAGGTATCCCCAGTGCTGGTTTTAGTCCCCTTCGTTTTTGGGGTAGTCTAAATCCCCTATAGTGAGTCGTATTAatttc |
| Nocardia farcinica | gaaatTAATACGACTCACTATAGGGCGCGAGGGCGAGGCCATCGCCTGGCA | CGGACGGATCGCCTCGTGTGCCTCCTG | ACTCGACCCGGACCTGGTCGACGCGCAGGTTTTAGTCCCCTTCGTTTTTGGGGTAGTCTAAATCCCCTATAGTGAGTCGTATTAatttc |
| Mycobacterium tuberculosis | gaaatTAATACGACTCACTATAGGGACCGTGAGGGCGAAGCTATTGCCTGGCA | GGGCCGGATAGCCTCGTGCGCTTCCTG | CCTCAAACCGCGCATACCGGTAAAGCGGGTTTTAGTCCCCTTCGTTTTTGGGGTAGTCTAAATCCCCTATAGTGAGTCGTATTAatttc |
| Anaplasma phagocytophilum | gaaatTAATACGACTCACTATAGGGGATCGCGAAGGAGAGGCGATAGCTTGGCA | GTTGGACGGATAGCTTCGTGCGCTTCCTG | AGATGTTACAGTGAACAGGATGGTGTTCGTTTTAGTCCCCTTCGTTTTTGGGGTAGTCTAAATCCCCTATAGTGAGTCGTATTAatttc |
| Aeromonas hydrophila | gaaatTAATACGACTCACTATAGGGGATAGAGAGGGAGAAGCGATCGCCTGGCA | GGGCGGATCGCCTCGTGGGCCTCCTG | CCAAGACCGCCATCCAGGAGGCGTTTGCGTTTTAGTCCCCTTCGTTTTTGGGGTAGTCTAAATCCCCTATAGTGAGTCGTATTAatttc |

| Table S3: Stain information for resistance panel and CARMEN v2 | | |
|---|---|---|
| **Strain label** | **Species** | **Source** |
| MSSA | Staphylococcus aureus | Reference strain |
| MRSA1 | Staphylococcus aureus | BWH clinical isolate |
| MRSA2 | Staphylococcus aureus | Barczak lab at MGH |
| CSE1 | Klebsiella pneumoniae | BWH clinical isolate |
| CSE2 | Klebsiella pneumoniae | BWH clinical isolate |
| CRE1 | Klebsiella pneumoniae | BIDMC clinical isolate |
| CRE2 | Klebsiella pneumoniae | |
| CRE3 | Klebsiella pneumoniae | |
| CRE4 | Klebsiella pneumoniae | |
| CRE5 | Klebsiella pneumoniae | |
| CRE6 | Escherichia coli | |
| CRE7 | Escherichia coli | MGH clinical isolate |
| CRE8 | Klebsiella pneumoniae | BWH clinical isolate |
| CRE9 | Escherichia coli | ATCC |
| CRE10 | Klebsiella pneumoniae | ATCC |
| CRE11 | Klebsiella pneumoniae | Wadsworth lab clinical isolate |
| CRE12 | Klebsiella pneumoniae | Wadsworth lab clinical isolate |
| CRE13 | Escherichia coli | Wadsworth lab clinical isolate |
| CRE14 | Escherichia coli | Wadsworth lab clinical isolate |
| CRE15 | Escherichia coli | Wadsworth lab clinical isolate |
| CRE16 | Klebsiella pneumoniae | MGH clinical isolate |
| CRE17 | Klebsiella pneumoniae | |
| VSE | Enterococcus faecalis | Barczak lab at MGH |
| VRE1 | Enterococcus faecium | Barczak lab at MGH |
| VRE2 | Enterococcus faecalis | BWH clinical isolate |
| VRE3 | Enterococcus faecalis | BWH clinical isolate |
| MSSA | Staphylococcus aureus | Reference strain |
| MRSA | Staphylococcus aureus | BWH clinical isolate |

**Table S2: Primer and guide sequences for resistance panel**

| Gene | Forward primer | Reverse primer | Guide (reverse complement with T7 promoter attached) |
|---|---|---|---|
| mecA | gaaatTAATACGACTCACTATAGGGTCAACAAGTTCCAGATTACAACTTCACCAG | ATCTGATGATTCTATTGCTTGTTTTAAGTCGATA | GCAATGATTGGGTTAAATAACAAAACATGTTTTAGTCCCCTTCGTTTTTGGGGTAGTCTAAATCCCCTATAGTGAGTCGTATTAatttc |
| mecC | gaaatTAATACGACTCACTATAGGGTCAACAAATTTCAAATCACTACATCACCAG | GTCTGATGATTCTATTGCTTGCTTTAAATCGATA | TCTATTATAGCCTTAAAAGAAAATAAACGTTTTAGTCCCCTTCGTTTTTGGGGTAGTCTAAATCCCCTATAGTGAGTCGTATTAatttc |
| vanA | gaaatTAATACGACTCACTATAGGGTACATTGGAATTACGAAATCTGGTGTATGG | ACTTGCCATGCAAAGCTGAAAATGCTACAT | GCACGGATTACTTGTTAAAAAGAACCATGTTTTAGTCCCCTTCGTTTTTGGGGTAGTCTAAATCCCCTATAGTGAGTCGTATTAatttc |
| vanB | gaaatTAATACGACTCACTATAGGGTACATCGGAATTACAAAAAACGGTGTATGG | ATTTGCCATGCAAAACCGGGAAAGCCACAT | GCATGGGCTGCTTGTCATGAAAGAAAGCGTTTTAGTCCCCTTCGTTTTTGGGGTAGTCTAAATCCCCTATAGTGAGTCGTATTAatttc |
| vanC | gaaatTAATACGACTCACTATAGGGACCATTGGCATCGCACCAACAATGGATTGG | ACTTCCCATGCAAGACTGGAAAGAGGACAT | TTCTAGCCAAGGATTTATATTAGGAGAAGTTTTAGTCCCCTTCGTTTTTGGGGTAGTCTAAATCCCCTATAGTGAGTCGTATTAatttc |
| vanE | gaaatTAATACGACTCACTATAGGGAAAATAGGGATCACCGAAGAAGGTCATTGG | AACCTCCATGTAAAACTGGGAATAAAATAT | CTGTGAAGAAATCGTAGTTGATTTCGCAGTTTTAGTCCCCTTCGTTTTTGGGGTAGTCTAAATCCCCTATAGTGAGTCGTATTAatttc |
| vanG | gaaatTAATACGACTCACTATAGGGCCAATAGGAATTACAAGAAGTGGTGAATGG | TTTTGCCATGCAATACGGGGAATACCAAAT | ATCTATGCCCTGTTGTCGTTTCCCAAAAGTTTTAGTCCCCTTCGTTTTTGGGGTAGTCTAAATCCCCTATAGTGAGTCGTATTAatttc |
| kpc | gaaatTAATACGACTCACTATAGGGCGTCTAGTTCTGCTGTCTTGTCTCTCATGG | CAAAGTCCTGTTCGAGTTTAGCGAATGGTT | GCTGGCTGGCTTTTCTGCCACCGCGCTGGTTTTAGTCCCCTTCGTTTTTGGGGTAGTCTAAATCCCCTATAGTGAGTCGTATTAatttc |
| ndm1 | gaaatTAATACGACTCACTATAGGGAATGTCTGGCAGCACACTTCCTATCTCGAC | TGATCTCCTGCTTGATCCAGTTGAGGATCT | CAACGGTTTGATCGTCAGGGATGGCGGCGTTTTAGTCCCCTTCGTTTTTGGGGTAGTCTAAATCCCCTATAGTGAGTCGTATTAatttc |
| oxa | gaaatTAATACGACTCACTATAGGGTTAAAATTCCCAATAGCTTGATCGCCCTCG | ATAAACAGGCACAACTGAATATTTCATCGC | CCAAGTCTTTAAGTGGGATGGACAGACGGTTTTAGTCCCCTTCGTTTTTGGGGTAGTCTAAATCCCCTATAGTGAGTCGTATTAatttc |
| vim | gaaatTAATACGACTCACTATAGGGGATGAGTTGCTTYTKATTGATACAGCKTGG | GATAGAARASYTCTACKGGACCGAARCGCA | CTCGCGGAGATTGAAAAGCAAATTGGACGTTTTAGTCCCCTTCGTTTTTGGGGTAGTCTAAATCCCCTATAGTGAGTCGTATTAatttc |
| imp | gaaatTAATACGACTCACTATAGGGAMAGATACTGAAAADTTAGTHAVTTGGTTT | CAGGYARCCAAACYACTASRTTATCTKGAG | GAATAGAGTGGCTTAATTCTCAATCTATGTTTTAGTCCCCTTCGTTTTTGGGGTAGTCTAAATCCCCTATAGTGAGTCGTATTAatttc |
| ctxm15 | gaaatTAATACGACTCACTATAGGGATAAAACCGGCAGCGGTGGCTATGG | GCTAATACATCGCGACGGCTTTCTGCCTTA | ATCGTGCGCCGCTGATTCTGGTCACTTAGTTTTAGTCCCCTTCGTTTTTGGGGTAGTCTAAATCCCCTATAGTGAGTCGTATTAatttc |
| mcr-1 | gaaatTAATACGACTCACTATAGGGGCTCGTTGGCTTAGATGACTTTGTCGCTGC | GCTTAAAATACGCAGGCCCGTGATTGCCCA | GGCAAAGATATGCTGATCATGCTGCACCGTTTTAGTCCCCTTCGTTTTTGGGGTAGTCTAAATCCCCTATAGTGAGTCGTATTAatttc |
